# VIsoQLR: an interactive tool for the detection, quantification and fine-tuning of isoforms using long-read sequencing

**DOI:** 10.1101/2022.03.01.482488

**Authors:** Gonzalo Núñez-Moreno, Alejandra Tamayo, Carolina Ruiz-Sánchez, Marta Cortón, Pablo Mínguez

## Abstract

DNA variants altering the pre-mRNA splicing process represent an underestimated cause of human genetic diseases. Their association with disease traits should be confirmed using functional assays from patient cell lines or other alternative models to detect the formation of aberrant mRNAs. Long-read sequencing is a suitable technique to identify and quantify mRNA isoforms. Available isoform clusterization and/or quantification tools are generally designed for the whole transcriptome analysis. Experiments focusing on a single locus analysis need more precise data fine-tuning and visualization tools.

Here we describe VIsoQLR, an interactive analyzer, viewer and editor for the semi-automated identification and quantification of known and novel isoforms using long-read sequencing data. VIsoQLR is tailored to thoroughly analyze mRNA expression and maturation in low-throughput splicing assays. This tool takes sequences aligned to a reference, defines consensus splice sites, and quantifies isoforms. Users can edit splice sites through dynamic and interactive graphics and tables as part of their manual curation. Known transcripts, or isoforms detected by other methods, can also be imported as references for comparison. Here, we explain VIsoQLR principles and features, and show its applicability in a case study example using Nanopore sequencing. VIsoQLR is available at https://github.com/TBLabFJD/VIsoQLR.

## INTRODUCTION

Spliceogenic DNA variants are an underestimated cause of genetic diseases (Lord & Baralle, 2021). They disrupt canonical donor and acceptor splicing sites or introduce cryptic exonic or intronic sites leading to aberrant mRNA maturation processes. Detection of genomic variation modifying splicing patterns and their association to disease traits is a challenging task that requires *in vivo* or *in vitro* functional studies. Global transcriptomics approaches are available (Mehmood et al., 2020); however, analysis is usually restricted to a single locus (or few loci) in Mendelian diseases. In these cases, some accurate and cost-effective procedures to detect altered expression in a targeted region are splicing RT-PCR assays applied to patient samples (Anna & Monika, 2018) or minigenes-based exon trapping assays if the damaged primary tissue is not accessible (Cooper, 2005).

Traditionally used analytical techniques have significant drawbacks in addressing the full spectrum of splicing events. First, Sanger sequencing or capillary electrophoresis assays are time-consuming techniques with laborious protocols that cannot assess the relative level of transcript isoforms. On the other hand, short-read sequencing of RT-PCR products cannot determine the exon organization of the full-length isoforms. The recent advent of long-read sequencing (LRS) appears as an alternative to characterize and quantify the isoform spectrum in splicing assays (Amarasinghe et al., 2020).

Several bioinformatics tools are available for the quantification and analysis of transcript isoforms. Most of them define isoforms as clusters of reads. In addition, they may require: 1) a reference sequence(s), such as StringTie2 (Kovaka et al., 2019); 2) the reference and an annotation file with exon coordinates, as is the case of Mandalorion (Byrne et al., 2017), TALON (Wyman et al., 2019), FLAIR (Tang et al., 2020) and LIQA (Hu et al., 2021); 3) a reference and short-read sequencing data to infer splice sites, such as FLAIR and IDP (Fu et al., 2018); or 4) none of the above, such as Oxford Nanopore algorithm (https://github.com/epi2me-labs/wf-transcriptomes). Pacific Biosciences’ (PacBio) sequencing data have its own collection of available tools, including PacBio IsoSeq3 (Gonzalez-Garay, 2016), IsoCon (Sahlin et al., 2018), SQANTI (Tardaguila et al., 2018), TAPIS (Abdel-Ghany et al., 2016), and SpliceHunter (Kuang & Canzar, 2018). To the best of our knowledge, there is no interactive tool that allows close inspection and fine-tuning analysis of one gene at a time on data from low-throughput approaches.

Herein, we present VIsoQLR, an interactive analyzer, viewer and editor for identifying and quantifying isoforms obtained from LRS data without prior knowledge of splice sites. VIsoQLR is designed to characterize aberrant mRNAs detected by functional assays targeting a single *locus* linked to specific phenotypes.

## MATERIALS AND METHODS

### Software implementation and availability

VIsoQLR is implemented in R (R Core Team, 2020) using the Shiny package (Chang et al., 2021) to build the interactive local web-based application. Figures displayed in the app are rendered using the plotly package (Sievert, 2020). VIsoQLR code, installation and user manual are available at https://github.com/TBLabFJD/VIsoQLR. A docker image can also be downloaded from https://hub.docker.com/r/tblabfjd/visoqlr.

### Minigene splicing assay and RT-PCR

We cloned the region from exon 5 to 7 of the gene *PAX6* (RefSeq transcript NM_000280.4), including ∼200bp intronic sequence on each side into the exon trapping expression pSPL3 vector. This construction was transfected in HEK-293T cells. Total RNA was isolated and retrotranscribed to cDNA using random hexamers, as previously described (Tarilonte et al., 2022). The obtained cDNA was further amplified using a primer pair that hybridized with the exons SD6 and SA2 from the expression vector.

### Long-read sequencing and analysis

The amplified PAX6 cDNA was sequenced on a MinION Mk1B device (Oxford Nanopore Technologies, ONT, UK) using a SpotOn Flow Cell (R9.4.1). Library preparation was carried out using the SQK-LSK109 sequencing kit (ONT) following the recommended protocol “Native barcoding amplicons” (version NBA_9093_v109_revF_12Nov2019). The library was sequenced until 5000 reads were obtained. Base-calling was performed using Guppy v5.0.16.

Reads were mapped using the GMAP aligner (Wu & Watanabe, 2005). GMAP parameters were: ‘-n1’ to avoid chimeric alignments, ‘--cross-species’ for a more sensitive search for canonical splicing, ‘--gff3-add-separators=0’ to prevent the “###” separator after each query sequence, and ‘-f 2’ to generate a GFF3 file containing exons coordinates. In the case study described here, VIsoQLR was applied using the automatic detection of splice sites with default parameters and filtering out truncated reads.

Data was also analyzed with StringTie2 (Kovaka et al., 2019) to detect splicing isoforms. In this analysis, we mapped reads using GMAP with the same parameters as the above, setting ‘-f samse’ to generate a SAM file, which was transformed into a BAM file, sorted, and indexed. StringTie2 was run using long reads mode (using ‘-L’ parameter). The detected and quantified isoforms were retrieved in a GTF file and visualized using VIsoQLR.

### Semi-quantitative capillary electrophoresis

PCR was performed as described above, but the reverse primer targeting SA2 was HEX-labelled. Fluorescent amplicons were run together with ROX1000 size standard (Asuragen, USA) under denaturing conditions in an ABI3130xl Genetic Analyzer (Thermo Fisher Scientific, USA). The results were analyzed with GeneMapper software (Thermo Fisher Scientific, USA).

## RESULTS

### Software scope and description

VIsoQLR has been developed to provide users with a graphical and interactive tool for isoform identification and quantification using LRS data generated by Nanopore or PacBio technologies. It gives a single-locus at a time analysis as the data exploration focuses on the graphical display of splice site distribution across the gene. VIsoQLR automatically detects splice site coordinates that can be fine-tuned according to the user’s expertise.

Figure 1 shows the workflow for isoform analysis using VIsoQLR. First, raw reads need to be aligned to a reference sequence. Here we suggest the GMAP aligner based on the performance of our own experience on Nanopore sequences. Still, other mappers, such as minimap2 (Li, 2018), can be used. Mapped data is uploaded to VIsoQLR. Using this input as GFF3 or BED6 format, VIsoQLR defines consensus exon coordinates (CECs) based on the frequency of the reads’ exon coordinates (start and end positions). Start and end positions are treated independently. Some facilities for selecting final CECs are provided: 1) a frequency filter, in which start and end positions above a configurable frequency are selected as candidate CECs (by default 3%); 2) a position window, in which the user can define a window, where other non-candidate CEC are merged (by default 5 nucleotides (nt) on both sides of each candidate CEC); and 3) a filter in which candidate CECs are merged into the most frequent one if they are closer than a given distance (by default 3 bases). In addition, VIsoQLR allows the user to change any parameter that defines CECs automatically and to add, delete and edit them manually. Known splice sites can also be uploaded in a file. Thus, once the consensus start and end positions are selected, the exons are defined accordingly, and reads with the same exons are considered the same isoform. The final isoform collection is defined, presented and quantified. Reads that do not fit into the delimited exons are discarded. Any change in exon coordinates makes isoforms redefined and quantified again on the fly.

**Figure 1.**
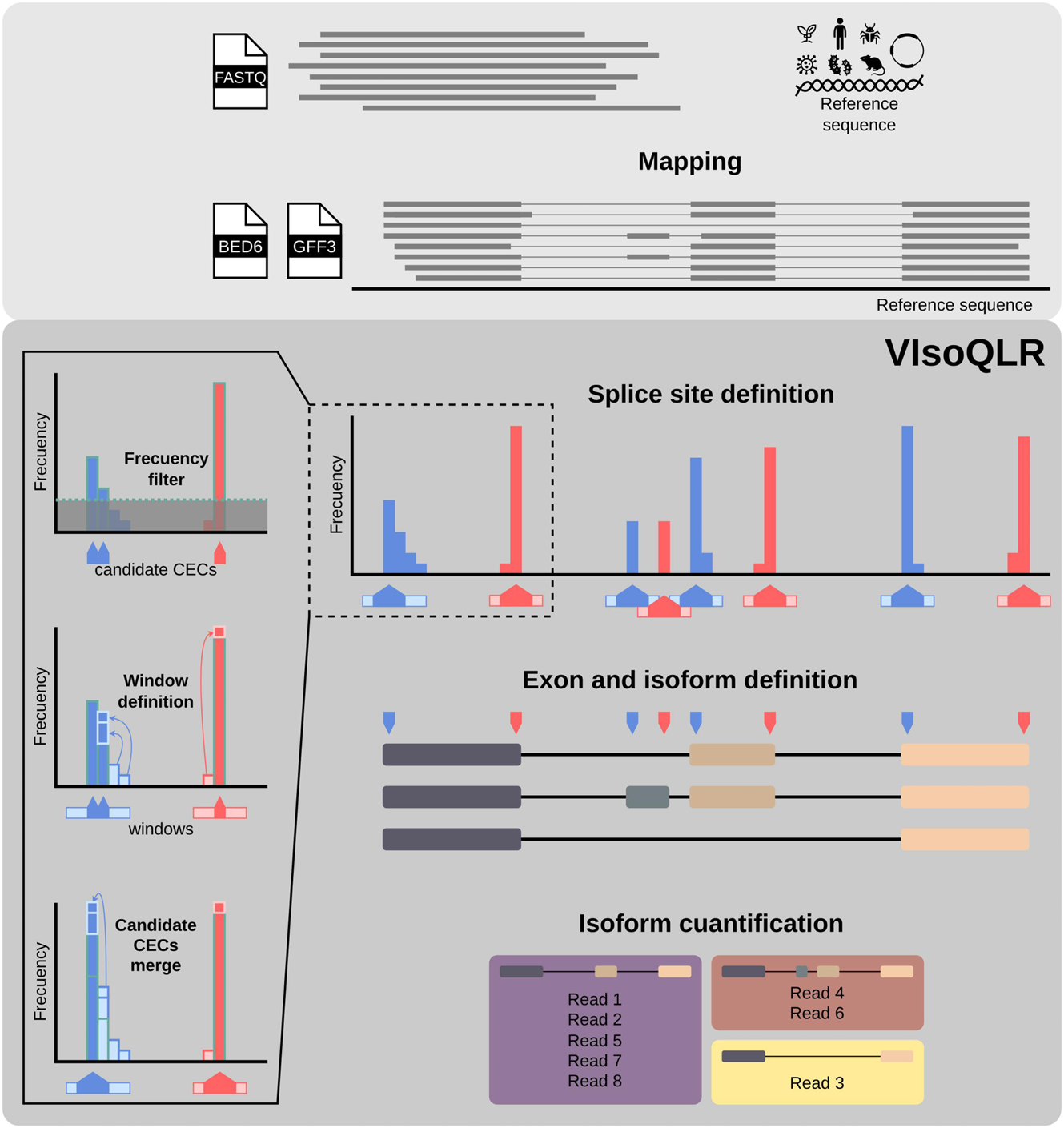
Workflow for isoform detection and quantification using VIsoQLR. Sequenced long-reads are mapped to generate a GFF3 or BED6 file containing the coordinates of all transcripts and exons. The frequency of each exon’s start and end coordinates are calculated with all reads. The selection of the consensus exon coordinates (CEC) includes the application of several optional features: 1) frequency threshold, which selects the most frequent ones; 2) window definition, where a window is defined around each candidate CECs so that close non-candidate CES are assigned to the nearest candidate CECs; 3) candidate CECs merge, where very close candidate CECs are merged into the most frequent one. Once the final CECs are defined, they are assigned to all read coordinates to define consensus exons and isoforms. Transcripts with all exons fully delimited by CECs are grouped into isoforms for quantification.

### User interface

The VIsoQLR control panel of the user interface is shown in Figure 2 (the complete VIsoQLR user interface is shown in Supplementary Figure 1). To start the analysis, the user must submit a GFF3 or BED6 file with aligned sequences at the top of the control panel (Figure 2A). Once reads are uploaded, the different sequences submitted are displayed in a drop-down menu in the analysis bounding section (Figure 2B). Sequences are analyzed one at a time. Also, the user can set the sequence boundaries from the same section to be explored. This is a useful option if the user wants to avoid working with exons coming from a vector or to focus on the splicing events of specific exons. Lastly, in this section, the user has the option to use only full-length transcripts. VIsoQLR has a default configuration to detect splice sites that can be changed in the “Exon coordinates” menu (Figure 2C). Here, options include different approaches to select CECs, such as setting up a minimum percentage of reads supporting the start and end exon coordinates, defining the size of the windows, or merging close CECs. In this control panel section, the user can upload previously defined exon coordinates (“Custom exon coordinates” menu) that replace or are merged with the automatically detected CECs performed by VIsoQLR.

**Figure 2.**
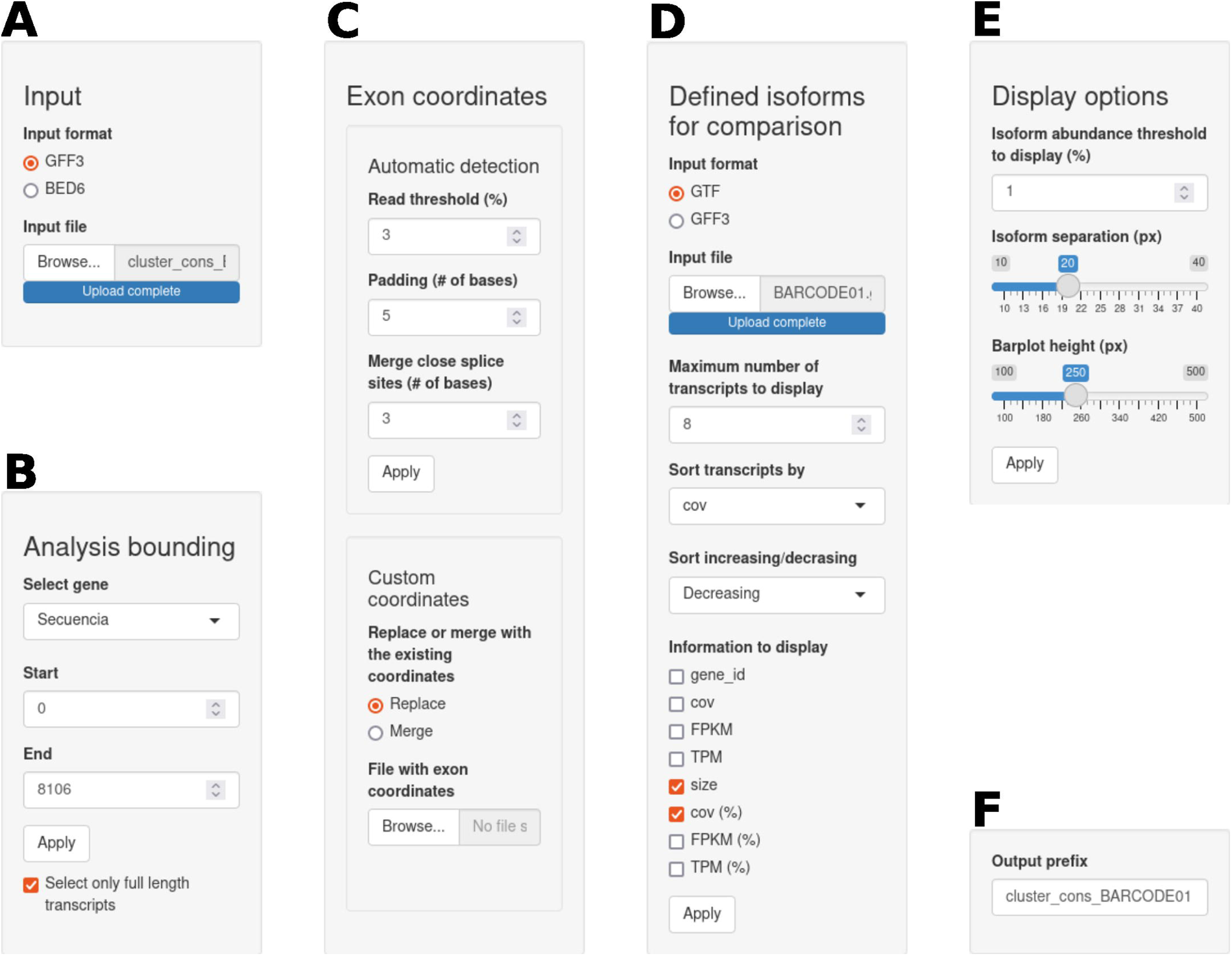
Control panel of the user interface of VIsoQLR. For visualization purposes, the control panel shown in the application as a single column is here split into six subpanels. **A**. Input subpanel where users can select the input file format and upload the aligned sequences. **B**. Analysis bounding subpanel allows the analysis of the gene and sequence area. It also contains an option to analyze only the full-length transcripts. **C**. Exon coordinates subpanel, with the option to automatically detect consensus exon coordinate (CEC). **D**. External isoforms subpanel, where users can upload known or previously defined transcripts as a reference to curate the isoforms detected by VIsoQLR. **E**. Display options subpanel. Here the user can filter the isoforms to be displayed based on their abundance and fine-tune the graphics. **F**. Download prefix subpanel is used to indicate the prefix of all downloadable tables and figures.

VIsoQLR also allows plotting known or previously analyzed transcripts for visual comparison with the isoforms detected in the experiment under analysis. They can be submitted in GTF, or GFF3 format (“Load transcripts from file” menu) (Figure 2D) and are displayed justified below isoforms with the same exon color codes to facilitate a direct comparison. Along with the transcript ID and size, it is possible to specify additional information about the transcript to be displayed. This information is stored in the last column of these two file formats. Visualizing a reference set of isoforms can help compare new results with previous data using VIsoQLR, those obtained with other software, or known transcripts.

The figure settings can be adapted to the user’s requirements using the “Display option” menu (Figure 2E). Lastly, all results, including figures and tables, can be downloaded by setting their prefix in the “Output prefix” menu (Figure 2F).

Figure 3A shows the collection of isoforms detected by VIsoQLR, including their exon configuration, coordinates, lengths and relative quantification. Here, isoforms detected by an external method using the same data uploaded by the user for reference or comparison purposes are shown. The exons of the “reference” isoforms are justified by those detected by VIsoQLR. Color coding is used to represent identical exons. Below, the start (in blue) and end (in red) exon sites are shown as peaks. The height of the peaks represents the percentage of reads mapped with the same position. A zoom and graphical selection of regions is also provided for close inspection of details. Selected CECs are marked with a dot, and when hovered cursor over, these dots display a tag with the exact coordinates and the percentage of reads supporting it. The figure can be downloaded as an HTML file maintaining all the dynamic properties and a static figure in many formats in the desired size.

**Figure 3.**
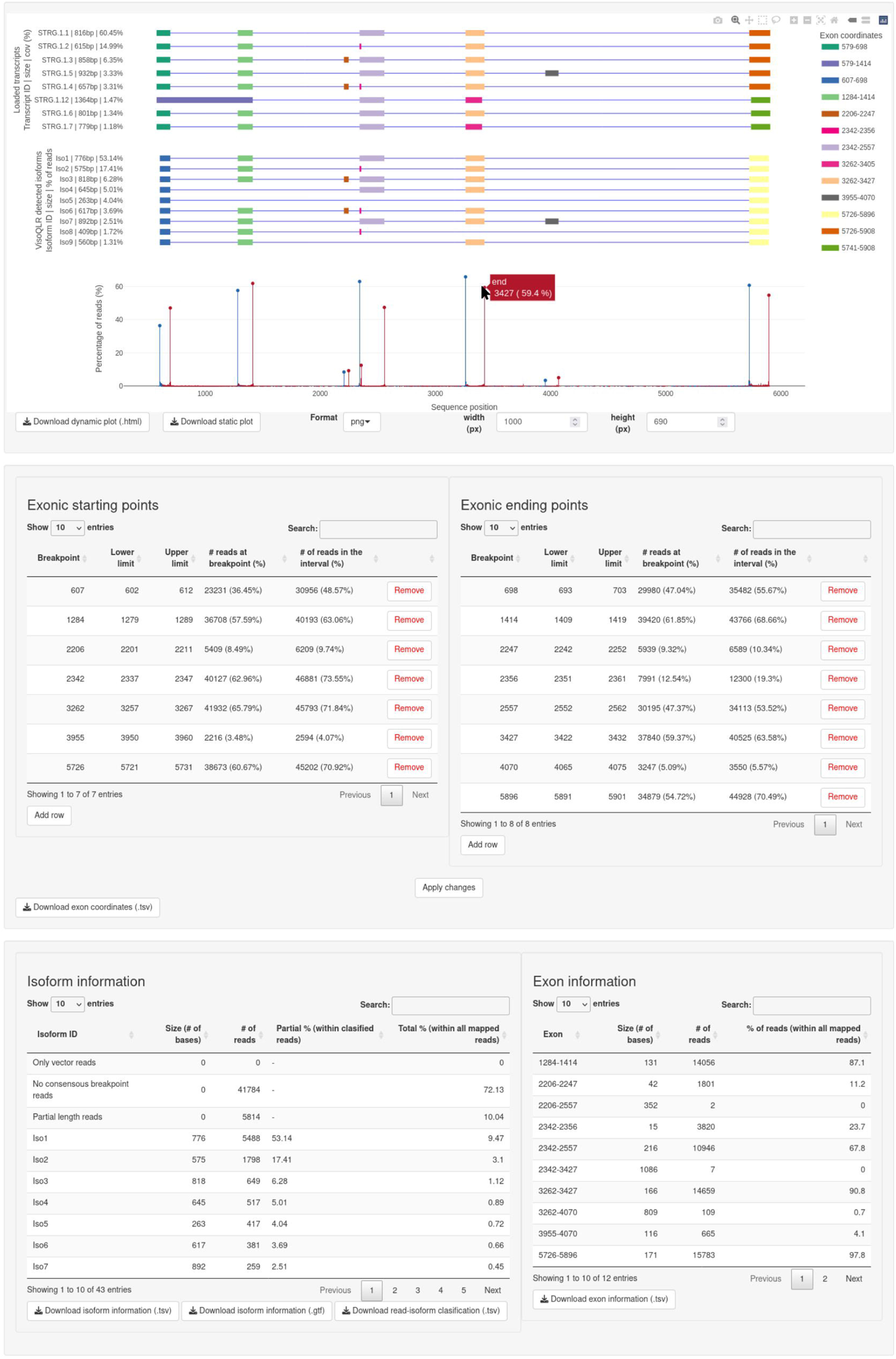
VIsoQLR results panel. **A**. Display subpanel containing the isoforms detected by VIsoQLR, including their exon configuration, coordinates, lengths and relative quantification. If uploaded by the user, externally defined isoforms are displayed. The color code is used to identify identical exons. Below isoforms, the frequency of start (blue) and end (red) coordinates are shown. The consensus exon coordinates (CECs) are marked with a dot on each bar, and the exact coordinate and frequency are displayed with the cursor over. All the plots are aligned on the x-axis. This plot can be downloaded as a dynamic figure in HTML or as a static figure in multiple formats in a configurable size. **B**. CECs are displayed in two tables (for start and end coordinates) with “Breakpoint”, “Lower limit”, and “Upper Limit” information that can be edited. The number of reads at the exact CECs and corresponding intervals are displayed. These coordinates can be downloaded as a single table. **C**. Extra isoform and exon information regarding their lengths and abundances is displayed and can be downloaded in multiple formats.

The consensus positions of the start and end exon are shown in two separate tables (Figure 3B). Their coordinates, as well as the window defining them, can be edited directly in these tables. Any edition made in these tables automatically redefines the exons and isoforms and recalculates their relative abundance. These tables provide additional information on the size and abundance of isoforms and exons (Figure 3C).

### Case study

To show the applicability of VIsoQLR, we performed a multi-exonic minigene assay for exons 5 to 7 of the *PAX6* gene using DNA from a healthy individual. The design of our cloned sequence in the exon trapping expression pSPL3 vector is depicted in Figure 4A. We sequenced the amplified cDNA on a MinION flow cell and mapped the generated reads with GMAP. The aligned reads were uploaded into VIsoQLR, and isoforms were calculated using default parameters and keeping only full-length transcripts. To have an external reference of the methodology to detect exons and characterize isoforms, we applied StringTie2 (Kovaka et al., 2019) to the same data (see Material and Methods). In Figure 4C, we show isoforms detected by VIsoQLR at the top, justified with the results obtained by StringTie2 that were uploaded using the “Load transcripts from file” menu. Both tracks show isoforms with an abundance above 1%, and they are sorted decreasingly by this field. The splicing isoforms from this sample were additionally analyzed by semi-quantitative capillary electrophoresis (CE) of fluorescent amplicons to estimate the proportion of each isoform and compare it with the results obtained by VIsoQLR. The fluorescent emission peaks of each isoform are represented in the electropherogram in Figure 4B.

**Figure 4.**
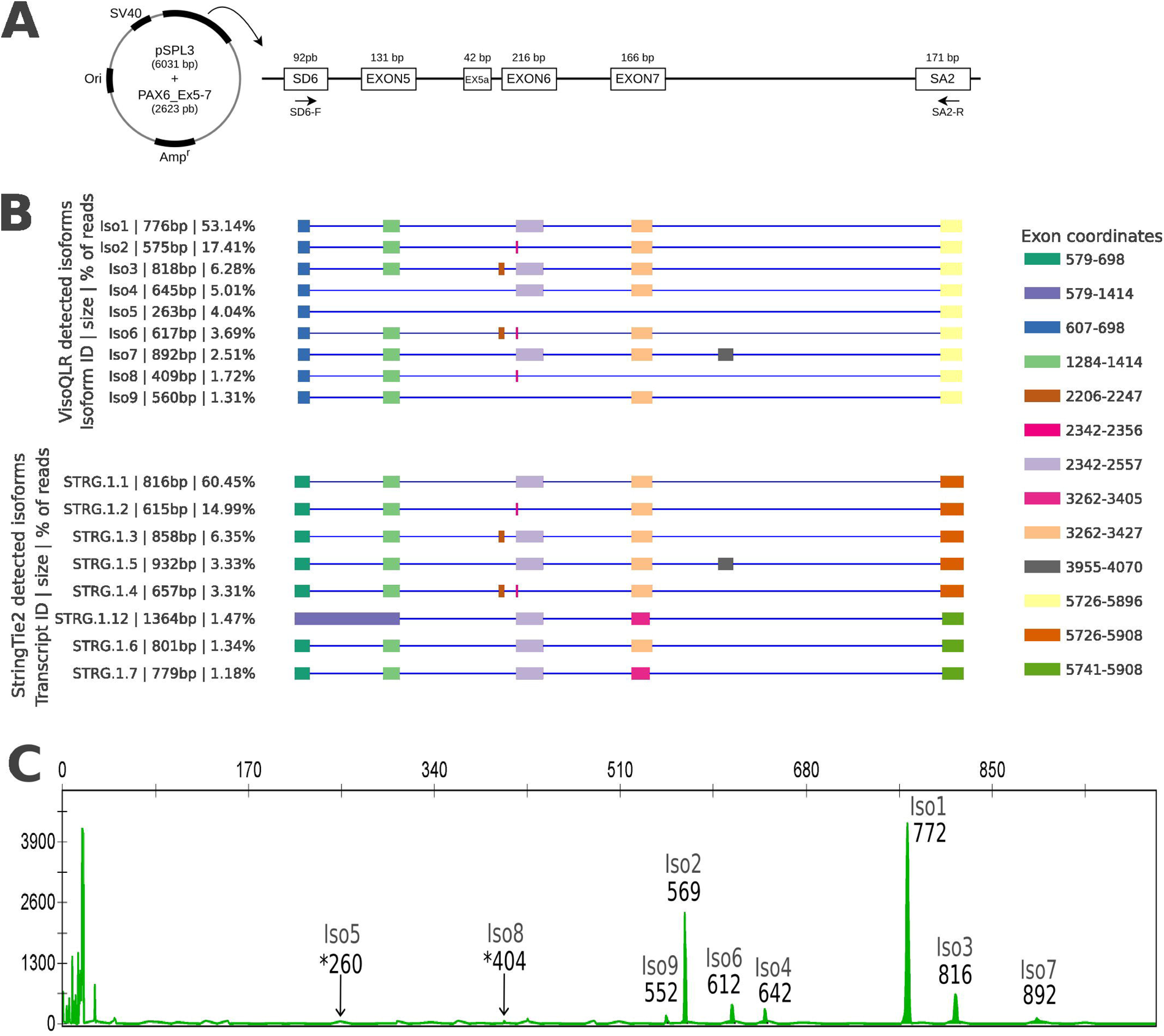
Minigene design and isoforms detected from the splicing assay. **A**. The exon trapping vector pSPL3 contains exons 5, alternative 5 (EX5a), 6, and 7 of *PAX6* (NM_000280.4). This vector contains a SV40 promoter, SD6 (splice donor 6) and SA2 (splice acceptor 2), and Ampicillin resistance gene (Amp^r^). The size in base pairs (bp) of the *PAX6* insert and the pSPL3 vector are shown. The forward (SD6-F) and reverse (SA2-R) primers are indicated. **B**. The isoforms detected by VIsoQLR are shown on the top track, including their exon configuration, coordinates, lengths and relative quantification. On the bottom track, isoforms detected by StringTie2 are shown. Only isoforms with an abundance above 1% are represented sorted by abundance. The color code is used to identify identical exons. **C**. Semi-quantitative PCR electropherogram. The x-axis represents the migration time, which correlates with the size of the molecules. The y-axis represents the absorbance intensity in Relative Fluorescence Units (RFU). The size, in base pairs (bp), is shown at the top of each peak. * Coordinates without well-defined peaks.

VIsoQLR detects 40 isoforms (Supplementary File 1), nine having an abundance above 1% of reads. In the case of StringTie2, it detects 58 isoforms (Supplementary File 1), eight of them composed of more than 1% of the reads. The most abundant isoform detected by VIsoQLR (53.1%) corresponds to the canonical transcript. This is composed of five exons (SD6-EXON5-EXON6-EXON7-SA2), including the two specific exons of the vector and three exons of *PAX6*, with a size of 776bp. StringTie2 also reports this as the most abundant isoform (60.5%), but different start and end positions make the transcript larger (816bp). In fact, all StrinTie2 isoforms have extra bases at the begging and end compared to VIsoQLR isoforms. This canonical isoform corresponds to CE’s highest peak at 772bp (Figure 4B).

The second most abundant isoform (17.4%) with 575bp detected by VIsoQLR is an alternative transcript in which the EXON6 was partially skipped due to the use of an alternative exonic donor site (Grønskov et al., 1999). This transcript also corresponds to the second most abundant isoform in StringTie2 (15.0 %) with a size of 615bp, and to the second highest emission peak at 569bp in CE. The third most abundant isoform called by both methods also coincides (6.3% and 6.4% in VIsoQLR and StringTie2, respectively). This is a second canonical transcript for *PAX6* in which an alternative exon 5a of 42bp (Figure 4A) is included (SD6-EXON5-EXON5a-EXON6-EXON7-SA2), making the isoform 818bp long in VIsoQLR and 858bp in StringTie2. This isoform also corresponds to the electropherogram’s third highest peak of 816bp.

Both methods also called some minor isoforms below 5%. Iso6 (3.6% and 617bp) and Iso7 (2.5% and 892bp) detected by VIsoQLR were also called by StringTie2, as STRG.1.4 (3.3% and 657bp) and STRG.1.5 (3.3% and 932bp), and correspond to in the electrophoretic peaks of 612bp and 892bp, respectively. However, four other minor isoforms were called by VIsoQLR but not by StringTie2. Iso4 (5.0% and 645bp) and Iso9 (1.3% and 560bp) appear in the electropherogram at 642bp and 552bp peaks, respectively. But, Iso5 and Iso8 were detected neither by StringTie2 nor CE. Finally, three isoforms (STRG.1.12, STRG.1.6, and STRG.1.7) were called only by StringTie2.

## DISCUSSION

Up to 15% of the pathogenic variants affect RNA splicing not only by involving canonical splicing sites but also by exonic and intronic non-canonical sites in increasing cases (Riolo et al., 2021). Therefore, they can be missed during conventional DNA screening or misinterpreted by *in silico* splicing predictions. Many algorithms have been developed to analyze and quantify the splicing pattern of genes using RNA sequencing (Byrne et al., 2017; Fu et al., 2018; Hu et al., 2021; Kovaka et al., 2019; Tang et al., 2020; Wyman et al., 2019). Although most can be used to analyze a single locus, they lack visualization and editing options that allow close exploration of regions of interest. VIsoQLR has been developed to fill this gap that is specifically required when performing low-throughput experiments. The study of Mendelian diseases is a clear example that requires this kind of facility. In their diagnosis, the seek for pathogenic spliceogenic variants usually is restricted to genes with a known association with the disease. In many of them, a single gene can explain most cases, e.g., *PAX6* and aniridia (Landsend et al., 2021), *ABCA4* and Stargardt disease (Cremers et al., 2020) or *NF1* and Neurofibromatosis type 1 (Koster et al., 2021). Functional characterization is crucial for correctly interpreting potential spliceogenic variants, especially in clinically actionable genes.

In addition to describing the VIsoQLR method and its features, we present a case study using LRS to illustrate its application. LRS is the state-of-the-art technology to study splicing (Amarasinghe et al., 2020). It allows the analysis of full transcripts, reaching sequences of about 10 Kb. It can report the whole repertoire of isoforms and provide their sequences. In contrast, classical approaches, such as capillary electrophoresis, only detect the most abundant isoforms and need a manual inspection of lengths to associate absorption peaks with known or *in silico* predicted isoforms.

Our case study aims to provide a comprehensive report of the *PAX6* isoforms in a healthy individual. Mutations in *PAX6* are responsible for nearly 100% of the cases of congenital aniridia, a rare developmental disease characterized by abnormalities in the iris and fovea (Blanco-Kelly et al., 2021). Two hotspot exons, EX5 and EX6, are prone to suffer naturally alternative splicing, resulting in a mixture of different splicing isoforms (Tarilonte et al., 2022). We compared results for a multi-exon *PAX6* minigene splicing assay provided by VIsoQLR using its automatic isoform detection with those detected by semi-quantitative CE. This comparison includes features not only specific to VIsoQLR but also those of LRS. VIsoQLR detected all isoforms present in the electropherogram, and there is a correlation in the abundance of the ranked isoforms provided by both methods. As expected, VIsoQLR provides a more extensive set of isoforms detected. However, some caveats in the abundance of small isoforms should be considered, as LRS tends to overrepresent them (Amarasinghe et al., 2020). An example of this might be the isoforms Iso5 and Iso8 detected by VIsoQLR (Supplementary Figure 2) with an abundance of 4% and 1.7%, respectively, but hardly distinguishable from the noise in the electropherogram (Figure4).

In addition, we also provide an example of how VIsoQLR can be used to visualize and compare results coming from other isoform detection algorithms. In this example, we chose StringTie2, a popular transcriptome analysis for short and long reads (Kovaka et al., 2019). The first evident difference is that StringTie2 reports a few extra bases for all isoforms at the outer boundary of the first and last exons compared to VIsoQLR. This extension seems to be an artifact as it is not present in BAM files used by StringTie2 nor in the GFF3 file used in VIsoQLR (Supplementary Figure 3 and Supplementary File 2). This can also be why StringTie2 reports systematically longer isoforms than electropherogram and VIsoQLR. In any case, the relative proportion of the most abundant isoforms detected by StingTie2 correlates with the isoforms reported with VIsoQLR and CE.

VIsoQLR demonstrates an accurate isoform detection using LRS data. On top of this, it provides a flexible, interactive and editable visualization framework for the manual inspection of potential gene splice sites. This feature allows a fast custom analysis of single-locus that is not available in any other tool. VIsoQLR can also be used to manually curate results from other isoform detection algorithms by adding its complete visualization and editing features. The docker containerization plus user interface allows users without deep knowledge in bioinformatics to analyze their data in a user-friendly program.

## Author contributions

GNM has conceived, designed and coded the tool and analyzed the data. MC, CRS and AT designed and performed experiments of the case study. GNM, PM and MC wrote the paper. PM and MC conceived the work and obtained the funding. All authors reviewed approved the manuscript.

## Data availability

VisoQLR code is available at https://github.com/TBLabFJD/VIsoQLR, docker image is available at https://hub.docker.com/r/tblabfjd/visoqlr. Case study data is available at https://github.com/TBLabFJD/VIsoQLR/example

